# Fine ultrastructural organization of glomerular neuropil in the developing zebrafish olfactory bulb

**DOI:** 10.64898/2026.09.18.752720

**Authors:** Ruth E. Montaño Crespo, Nila R. Mönig, Johannes Kappel, Alexandra Graff Meyer, Tomáš Gancarčik, Michal Januszewski, Rainer W. Friedrich

## Abstract

Across animal phyla, odor information is represented in the first olfactory processing center by combinatorial activation of discrete glomeruli. To examine how this organization of olfactory processing channels emerges during development we combined 2-photon calcium imaging, volume electron microscopy, and large-scale automated neuron reconstruction in zebrafish larvae. We focused on a developmental stage (130 hours post fertilization) when the olfactory bulb (OB) is structured into ∼16 large neuropil regions referred to as protoglomeruli but most glomeruli are not yet differentiated. Our ultrastructural reconstructions revealed that protoglomeruli are organized into smaller units referred to as microglomeruli, each of which is defined by a distinct set of mitral cells with overlapping, spatially confined dendritic arbors. Neighboring microglomeruli were innervated by distinct subsets of sensory axons and activity was more highly correlated among mitral cells associated with the same microglomerulus than across microglomeruli of the same or different protoglomeruli. These results reveal a precise subcellular organization of the neuropil in the developing OB into distinct processing channels, indicating that the layout of the adult glomerular array is established already early in development.

## Introduction

The olfactory system generates representations of complex, variable, and high-dimensional molecular stimuli to extract relevant information about the environment and to inform behavior. In vertebrates and invertebrates, individual olfactory sensory neurons (OSNs) expressing the same chemosensory receptor converge onto discrete glomeruli in the first central processing center, the olfactory bulb (OB) or antennal lobe, respectively (Buck, 2000; Wilson and Mainen, 2006). Olfactory sensory input to the brain is therefore organized into a map of discrete input channels, each representing a defined receptor. Within glomeruli, OSN axons make glutamatergic synapses onto principal neurons and subsets of inhibitory interneurons. Odors evoke specific combinatorial patterns of activity across the array of glomeruli that are reorganized by neuronal circuits containing prominent inhibitory connectivity motifs. These circuits normalize and decorrelate activity patterns evoked by different odors, resulting in a whitening of activity patterns that supports odor classification and association (Wilson and Mainen, 2006; Friedrich, 2013; Wanner and Friedrich, 2020).

In the mammalian OB, glomerular development comprises multiple sequential events including the initial guidance of sensory axons towards their target region and the subsequent convergence onto discrete glomeruli. These processes are mediated by a combination of molecular and activity-dependent axon guidance and sorting mechanisms in OSNs that are largely independent of their postsynaptic targets (Treloar et al., 1999; Potter et al., 2001; Sakano, 2010, 2020). Shortly after glomerular convergence, dendrites of immature principal neurons, the mitral cells (MCs), are recruited into the glomerular neuropil (Malun and Brunjes, 1996; Treloar et al., 1999; Matsutani and Yamamoto, 2000; Blanchart et al., 2006; Nishizumi et al., 2019). Subsequently, axonal targeting is refined (Zou et al., 2004) and primary dendritic arbors of MCs become restricted to individual glomeruli (Malun and Brunjes, 1996; Matsutani and Yamamoto, 2000; Blanchart et al., 2006; Fujimoto et al., 2023; Imai, 2025). As a consequence, canonical mitral cells of the adult OB receive direct sensory input through a single glomerular channel representing a defined odorant receptor. However, the developmental processes underlying the formation of discrete glomerular processing channels are not fully understood. In rodents, for example, some studies indicate that the array of glomeruli emerges within a few days after birth (Malun and Brunjes, 1996; Matsutani and Yamamoto, 2000; Blanchart et al., 2006; Nishizumi et al., 2019) while others reported the continuous addition of new glomeruli over weeks of postnatal development (LaMantia and Purves, 1989; Pomeroy et al., 1990). Moreover, as glomerular development has been studied primarily in rodents, it remains unclear whether the underlying processes are conserved across vertebrate classes.

We examined developing glomeruli in zebrafish, which provide important advantages for structural and functional analyses of neuronal circuits (Friedrich and Wanner, 2021). The OB of adult zebrafish contains a stereotyped glomerular map consisting of ∼140 anatomically distinct glomeruli (Baier and Korsching, 1994; Byrd and Brunjes, 1995; Braubach et al., 2012; Braubach and Croll, 2021) and additional densely packed glomeruli in a ventro-lateral region that cannot be delineated by conventional light microscopy (Mönig et al., 2026). At larval stages, in contrast, the OB contains a substantially smaller number of distinct and stereotyped neuropil domains referred to as protoglomeruli (Dynes and Ngai, 1998; Li et al., 2005, 2005; Braubach et al., 2013; Miyasaka et al., 2013; Shao et al., 2017). Zebrafish OSNs express different types of chemosensory receptors comprising conventional odorant receptors, trace amine-associated receptors (TAARs), vomeronasal type 2-like receptors (V2Rs), and a small number of vomeronasal type 1-like receptors (V1Rs) (Yoshihara, 2009; Manzini and Korsching, 2011; Miyasaka et al., 2013; Saraiva et al., 2015). The majority of these receptors are already expressed in an exclusive fashion at larval stages (Barth et al., 1996; Weth et al., 1996; Argo et al., 2003; Miyasaka et al., 2013; Shao et al., 2017), implying that individual protoglomeruli are innervated by multiple types of OSNs. As development proceeds, some protoglomeruli persist and grow only in size while others split into multiple smaller neuropil units (Li et al., 2005; Braubach et al., 2013). As a result, most glomeruli in the adult OB differentiate by segregation from protoglomerular precursors over a period of weeks. Throughout this period the neuronal circuitry in the zebrafish OB is already functional (Li et al., 2005; Mack-Bucher et al., 2007) and odors elicit innate behavioral responses (Lindsay and Vogt, 2004; Vitebsky et al., 2005; Whitlock, 2006; Mayseless et al., 2026), raising the question how stable function is maintained as the array of processing channels is dynamically reorganized during development.

Anatomical studies (Baier and Korsching, 1994; Braubach et al., 2012), activity measurements (Li et al., 2005), tracing of projections to higher brain areas (Miyasaka et al., 2014), and mapping of transcriptionally defined neuron types (Mayseless et al., 2026) revealed a non-random spatial organization of larval protoglomeruli that may support innate brain functions. Further studies provided insights into the molecular mechanisms that direct sensory axons to protoglomeruli in larval zebrafish (Shao et al., 2017; Cheng et al., 2022; Barnes et al., 2026). However, the organization of neurons within protoglomeruli remains poorly understood. Specifically, it remains unclear whether protoglomeruli are unstructured neuropil domains where individual mitral cells receive input from multiple types of convergent OSNs, or whether protoglomeruli exhibit an intrinsic organization that links OSN types to subsets of mitral cells prior to the differentiation of glomeruli.

The detailed anatomical analysis of the protoglomerular neuropil requires dense reconstructions of neurites from populations of neurons at ultrastructural resolution. We used volume electron microscopy (EM) (Denk and Horstmann, 2004) and automated segmentation by flood-filling networks (Januszewski et al., 2018) to reconstruct mitral cells in both OBs of a zebrafish larva after calcium imaging of odor-evoked activity. We found that mitral cells are organized within protoglomeruli into small clusters referred to as microglomeruli. Different microglomeruli contained dendrites of non-overlapping subsets of mitral cells that were innervated by specific sets of OSN axons. Activity correlations between mitral cells were significantly higher within than across microglomeruli. Further observations revealed midline-crossing dendritic arborizations of mitral cells and other neurons associated with a specific glomerulus. These results reveal a sub-protoglomerular organization of the OB neuropil into anatomically and functionally distinct microglomeruli, demonstrating that sensory processing channels are established with precision before glomeruli are fully differentiated. This organization may be the basis for the gradual segregation of glomeruli from protoglomerular precursors and for maintaining coherence of odor processing through development.

## Results

### Imaging odor-evoked activity in the larval zebrafish olfactory bulb

To obtain corresponding functional and ultrastructural data from the same neurons in the larval zebrafish OB we first measured odor-evoked activity in the OBs by two-photon calcium imaging and subsequently acquired a stack of EM images from the same specimen. Activity was measured in agarose-embedded, paralyzed Tg(alphatubulin:GCaMP6s, gad1b:DsRed) larvae at 130 hours post fertilization, which expressed the calcium indicator GCaMP6s throughout the brain (Diaz-Verdugo et al., 2019) and the red-fluorescent protein DsRed in most GABAergic neurons (Satou et al., 2013). Image series covering both OBs were acquired near-simultaneously from six optical planes, spaced by ∼14 μm, at a volume rate of 7.5 Hz (Fig. 1A-F) (Rupprecht et al., 2016). After completion of activity measurements, a high-resolution image stack (“anatomical reference”) was acquired before the sample was fixed and prepared for EM (Fig. 1G). Activity planes were slightly tilted relative to the anatomical reference (Fig. 1H) as a consequence of the remote scanning procedure used for fast volumetric imaging (Rupprecht et al., 2016). Responses were measured to six odors (two amino acids [Alanine, Serine], three bile acids [TDCA, TCA, GDCA] and cadaverine) and one control stimulus (E3 medium). Stimuli were applied for 5s at an inter-stimulus interval of 2 min and repeated three times in a pseudo-randomized block sequence (Methods).

**Figure 1:**
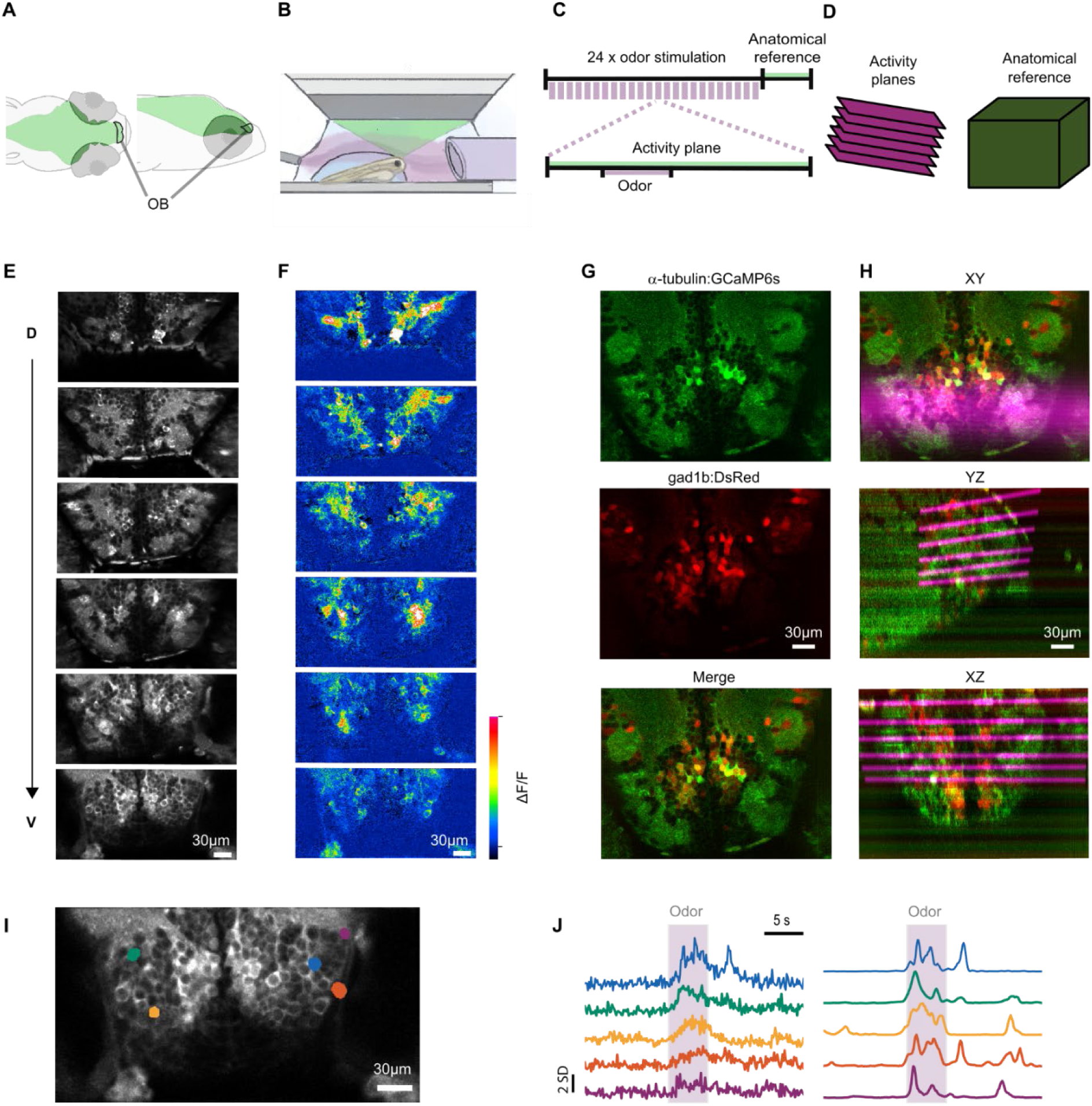
Imaging of odor-evoked activity in the larval zebrafish olfactory bulb. **A.** Olfactory bulb (OB) in larval zebrafish. **B.** Odor stimulation in a paralyzed, agarose-embedded larva under the objective. The larva was constantly superfused with medium through a frontal delivery tube and a caudal suction tube. Agarose in front of the nostrils was removed. **C.** Experimental schedule. Light purple indicates odor stimulation; green indicates image acquisition. The experiment comprised 24 pseudo-randomized odor trials followed by acquisition of the final anatomical reference. An odor trial consisted of a 6.7 s baseline acquisition, a 5 s odor stimulus pulse, and a 10 s post-stimulus period. **D.** Schematic: activity planes were slightly tilted relative to the anatomical reference. **E.** GCaMP6s expression (time-averaged images from six activity planes acquired during odor stimulation, ordered from dorsal to ventral). **F.** Relative change in fluorescence (ΔF/F) evoked by presentation of alanine in the same activity planes as in E. **G.** Slices through the anatomical reference showing pan-neuronal expression of α-tubulin:GCaMP6s (green) and Gad1b:DsRed expression in putative GABAergic neurons (red). **H.** Aligned activity planes superimposed onto the anatomical reference, shown in orthogonal views (top: xy, middle: yz, bottom: xz). Magenta regions represent activity planes convolved with a kernel representing the point spread function of the microscope in the z direction. **I.** ROIs representing five cells in the most ventral activity plane. **J.** ΔF/F traces (top) and inferred spike rates (bottom) for the ROIs shown in I, colors corresponding. Odor application indicated by gray shading. ΔF/F scale bar denotes two standard deviations (SD) of the baseline fluorescence. Inferred spike traces are normalized to the peak amplitude of each trace.

EM samples were prepared and screened after activity imaging from four zebrafish larva. One larva was then chosen for further analysis based on the quality of the EM image stack. In the six activity planes, regions of interest (ROIs) representing 701 neuronal somata were defined using Cellpose (Stringer et al., 2021) followed by manual proofreading (Fig. 1I). For each ROI, firing rates were inferred from time series of relative fluorescence changes (ΔF/F traces) using CASCADE (Rupprecht et al., 2021) in each trial (Fig. 1J). Putative GABAergic neurons were identified based on the DsRed signal (Fig. 1G).

### High-resolution structural acquisition and volume reconstruction

Volumetric EM data were acquired by serial block-face scanning EM (Denk and Horstmann, 2004; Denk et al., 2012). Sample blocks were prepared immediately after calcium imaging (Fig. 2A) using a modified fBROPA protocol (Genoud et al., 2018) and embedded in conductive silver resin (Wanner et al., 2016) (Fig. 2A). An image stack containing both OBs and part of the adjacent telencephalon (231 × 245 × 115 μm^3^) was acquired at a voxel size of 10 x 10 x 25 nm^3^ (Fig. 2B-D; Supplementary Fig. 1A,B). Individual images (tiles) were stitched and aligned into a coherent volume using a pipeline based on SOFIMA (Januszewski et al., 2024; Gancarcik and Hu, 2022) (Methods). In the resulting EM volume, ultrastructural details including fine processes, vesicles and other organelles could be clearly distinguished in all orientations (Fig. 2D).

**Figure 2:**
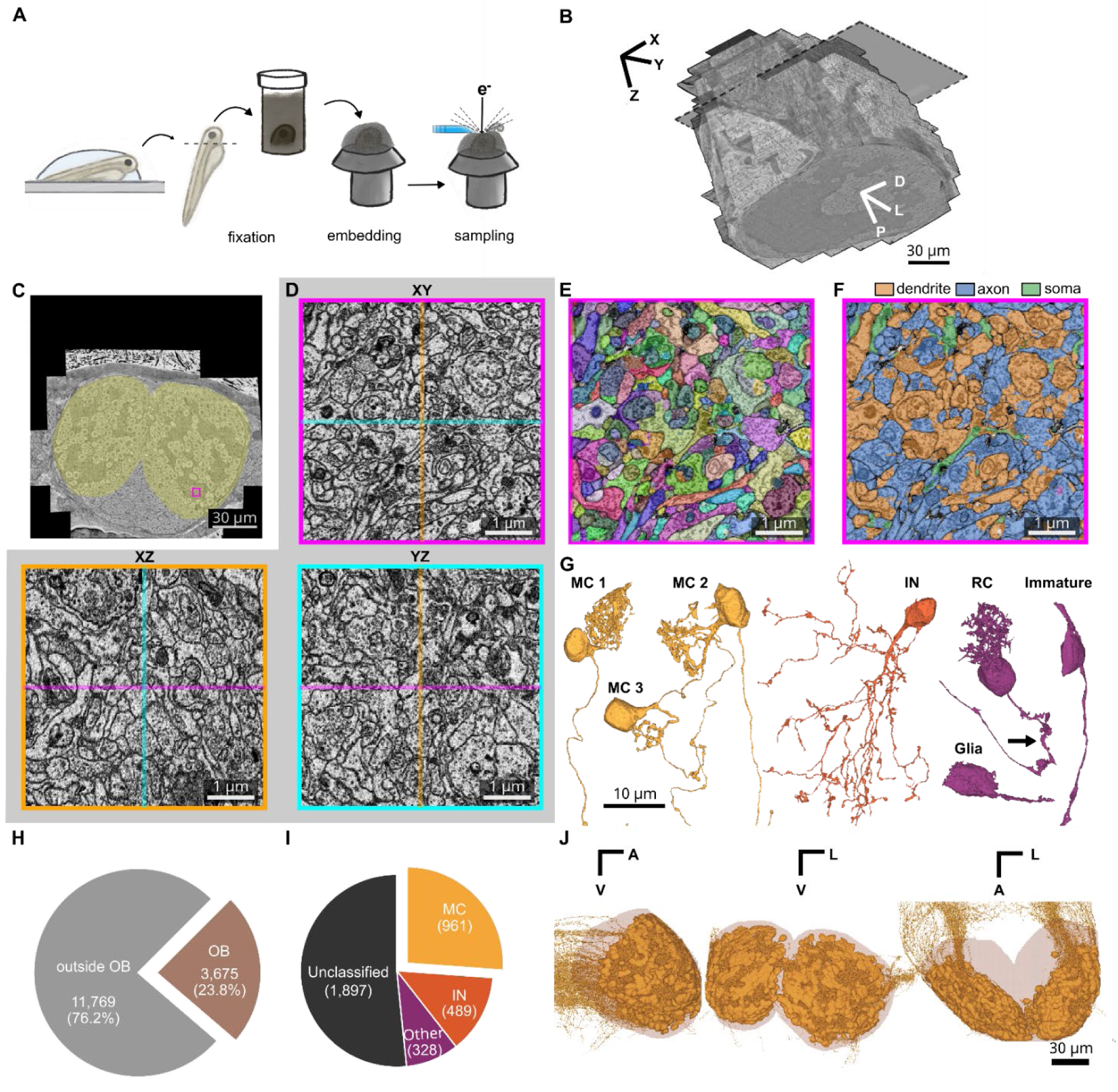
Volume electron microscopy and neuron reconstruction. **A.** Schematic of EM sample preparation following odor stimulation and calcium imaging. The head was fixed and processed using the fBROPA protocol and embedded in conductive resin for image acquisition by serial block-face scanning EM. **B.** EM volume (231 × 245 × 115 μm^3^) covering the OB and surrounding areas. Gray slice indicates the transverse plane shown in C. **C.** Transverse plane showing both OBs (yellow mask) along with surrounding tissue. Magenta square marks a representative neuropil-rich region enlarged in D. **D.** Orthogonal cross-sections through a neuropil-rich region at native resolution resolution (10 x 10 x 25 nm^3^; square in C). **E.** Base segmentation labels for the region in D. Different agglomerates are labeled using a randomized colormap. **F.** Semantic segmentation of the region shown in D. **G.** Representative cells of different types, colored by category: mitral cells (MC), interneurons (IN), and other. The “other” category includes ruffed cells (RC), immature neurons, and glia. Arrow depicts ruff. **H.** Number/fraction of nuclei in the EM volume located outside versus inside the OB. **I.** Number of nuclei inside the OB classified by cell type. **J.** Overlay of MCs, viewed in orthogonal projections. Note distribution of somata throughout the OBs and axonal projections leaving the OB.

Image data were segmented by flood-filling networks (Januszewski et al., 2018) using a model that was transferred from a previously trained model via the SECGAN domain adaptation algorithm (Januszewski and Jain, 2019). The resulting segments (Fig. 2E) contained very few false mergers, as evaluated by visual inspection, and were further semantically classified into dendrites, axons, nuclei and somata (Fig. 2F). Segments were then agglomerated by pretrained artificial networks (Januszewski et al., 2018) into larger structures that typically represented large parts of individual neurons (Fig. 2G).

3,675 (23.8%) of the 15,444 nuclei in the EM volume were located within the OBs (Fig. 2H). Agglomerates linked to these nuclei were inspected visually and selected for further analysis when they contained extensive neurites and allowed for morphological cell type classification. Discarded agglomerates were often small and included incomplete reconstructions that did not provide sufficient information to assess cell type identity. A substantial fraction of these agglomerates probably represented immature neurons, which were found predominantly in the medio-ventral OB and lacked input or output synapses. The set of selected reconstructions comprised 1,778 cells (48.4% of all nuclei in the two OBs; Fig. 2I) that were distributed approximately equally between the two OBs.

Reconstructed cells were assigned to one of three classes (Fig. 2G) by morphological criteria (Wanner et al., 2016): (1) MCs were distinguished by their characteristic compact dendritic tuft and soma position; (2) INs were identified by their more diffuse and spatially extended dendritic arbor and by the absence of an axon projecting to the telencephalon; (3) “Other” cells were defined as cells that did not match morphological criteria for MCs or INs. This category included immature neurons that lacked synapses and glia, many of which were found on the surface of the OB (Fig. 2G). In addition, it included 48 ruffed cells, a distinct type of OB output neuron in teleosts that is characterized by dense protrusions (the “ruff”) at the axon initial segment (Kosaka and Hama, 1979) (Fig. 2G; Supplementary Fig. 1E). To our knowledge, ruffed cells have previously been described only at late developmental stages (Fuller and Byrd, 2005; Mönig et al., 2026), possibly because small protrusions are difficult to detect by light microscopy (LM). In total, 961 cells were classified as MCs, 489 as INs, and 328 as other cells.

Obvious reconstruction errors of selected cells encountered during the initial inspection were corrected by repairing false splits or mergers. In addition, more detailed proofreading of MCs was performed in 136 of 961 MCs that appeared potentially incomplete. Edits made during proofreading primarily corrected split errors in the dendritic arbor. In MCs that underwent editing, the average fraction of voxels classified as dendritic was significantly lower than in unedited MCs before editing but this difference was eliminated after editing (Supplementary Fig. 1C, D). These observations indicate that targeted proofreading selectively enhanced the completeness of dendritic reconstructions in MCs, which are the anatomical basis for further analyses. A previous comprehensive inventory of the zebrafish OB at ∼108 hpf classified 745 neurons as MCs (Wanner et al., 2016). This number presumably includes ruffed cells because previous reconstructions did not identify protrusions at the axon initial segment. We therefore estimate that our dataset contains approximately two thirds of all MCs in each OB.

### Cross-modal registration of light microscopic and EM datasets

To map neurons analyzed in activity measurements to the EM dataset we registered the in vivo activity planes, the anatomical reference, and the EM volume from the same larva in two sequential steps (Fig. 3A). Activity planes were first registered within each trial by computing a similarity transform from a reference trial and applying it to all remaining trials of that plane (Methods). Activity planes were then aligned to the anatomical reference using tiled template matching. The EM volume was coarsely aligned to the anatomical reference by rotating and reslicing to match its dorsal view. Fine alignment was then performed in BigWarp (Bogovic et al., 2016) using two sets of landmarks in succession: a sparse set of coarse anatomical landmarks (blood vessels, identifiable neuropil regions), followed by a denser set derived from segmented EM nuclei matched to their corresponding cells in the anatomical reference. These nuclear matches were used to compute a thin-plate spline transform, which was iteratively refined as additional matches were added (Methods). This two-stage procedure enabled bidirectional mapping of coordinates and ROIs between the activity planes, the anatomical reference and the EM volume, allowing functional and structural data to be queried in a common space (Fig. 3A-C).

**Figure 3:**
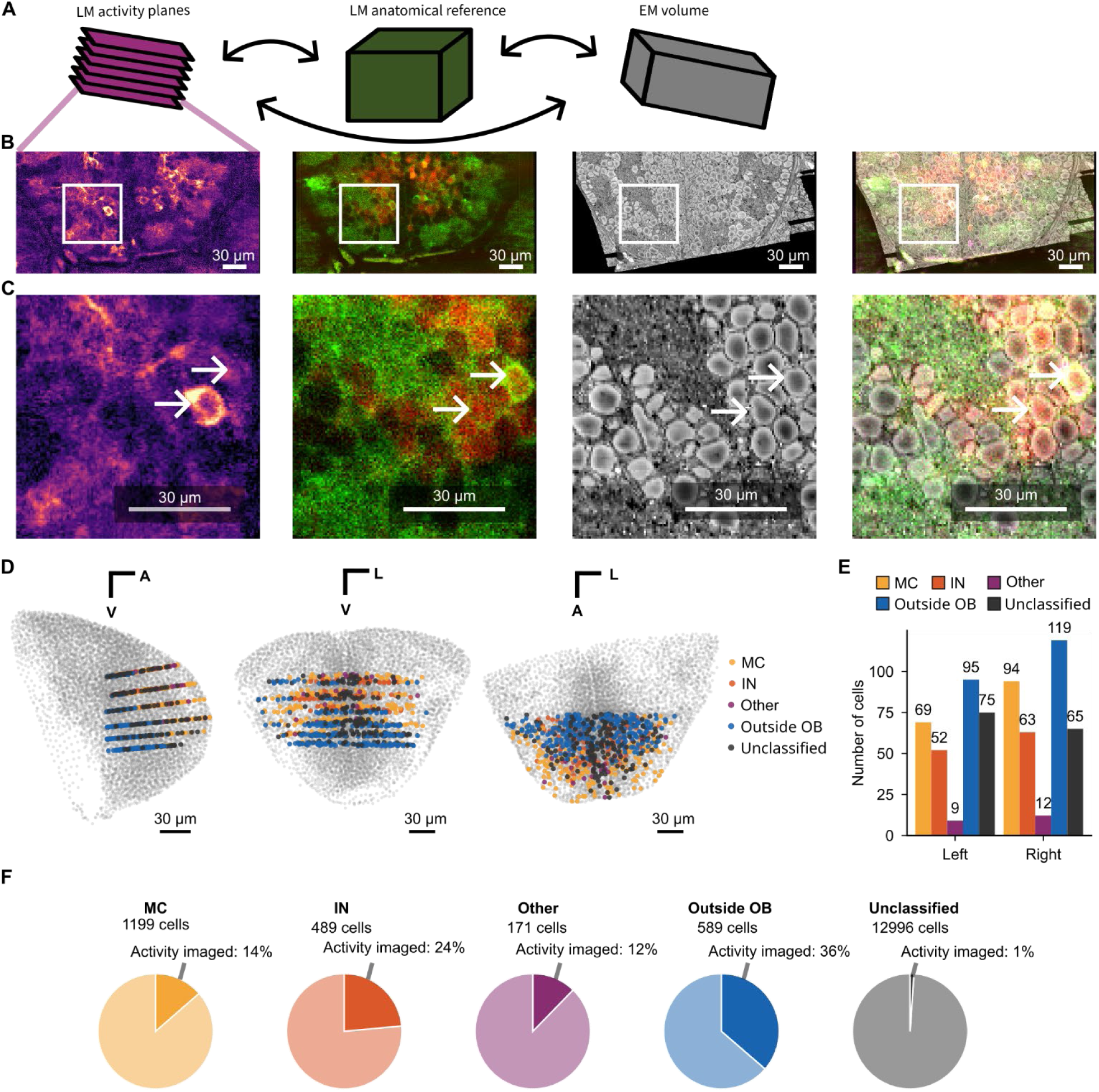
Cross-modal registration of functional and structural datasets. **A.** Schematic: cross-modal registration between activity planes (LM; slightly tilted relative to the optical axis), the anatomical reference (LM; perpendicular to the optical axis), and the EM volume. **B.** Representative images across modalities, shown in LM activity plane space. From left to right: reference activity plane, warped anatomical reference, warped EM volume, and overlay. **C.** Zoomed-in view of the white square in B across modalities as in B. White arrows depict examples of matched cells. **D.** Registered EM-segmented somata and LM activity plane ROI centroids, projected in different directions. ROI centroids are color-coded by cell type. **E.** Number of cells by cell type in the left and right OB. **F.** Percentage of cells in activity planes relative to all cells in the EM volume by cell type.

After registration, individual cells could be identified across all three datasets (Fig. 3B,C). To quantify the accuracy of LM-EM alignment we evaluated correspondences between somatic centroids using three metrics: mean alignment error (the Euclidean distance between each transformed LM centroid and its nearest-neighbor match in the EM point cloud), neighbor distance ratio (NDR; the nearest-neighbor distance normalized by the mean distance to 10 subsequent nearest neighbors) and a separation score (the difference between the residual distance to the assigned match and the distance to the nearest competing candidate) (Bae et al., 2025). We then examined how each metric varied under systematic small modifications of rotation, scale, and shift applied to the LM centroids (Supplementary Fig. 2G,I,K). Our mapping exhibited only a small (∼1 μm) translational shift relative to the optimum along the dorso-ventral axis that had minimal effects on accuracy while rotation and scale parameters were optimal. Distributions of all three metrics confirmed that the large majority of centroids were well aligned with sub-nuclear accuracy without an obvious spatial bias (Supplementary Fig. 2A-F).

Alignment of somata may be more challenging in regions where somata are densely packed. We therefore examined each alignment metric as a function of local soma density in the EM dataset (Supplementary Fig. 2H,J,L). Outlier points with poor alignment scores were distributed largely independently of local density across all three measures. Manual inspection of the lowest-scoring centroids showed that genuine misalignments were predominantly located outside the OB, presumably because landmarks for alignment were concentrated within the OB. Using manual inspection focusing on the centroids with lowest alignment scores within the OB we found that these centroids were matched correctly between datasets. Consistent results were obtained using all three metrics.

A total of 653 cells were matched between activity planes and the EM dataset, with similar cell type contributions in the left and right OB (Fig. 3D,E). While the EM volume covered both OBs in their entirety, activity planes sampled only approximately 10% of the OB volume, targeting regions that contain mature, odor-responsive neurons (Fig. 3D-F). Most unclassified neurons were located outside this volume, consistent with the assumption that many unclassified cells are immature neurons (Fig. 3D-F).

### Discrete structural subcompartments within protoglomeruli

We next examined the organization of MC dendrites within protoglomeruli. In each OB, 16 protoglomeruli were manually defined by outlining neuropil regions in 3D. The arrangement of protoglomeruli in the two OBs was approximately symmetrical and consistent with previous protoglomerular maps (Dynes and Ngai, 1998; Li et al., 2005; Braubach et al., 2013; Wanner et al., 2016; Shao et al., 2017). Individual protoglomeruli were identified based on their shape and position, and named following an established nomenclature (Braubach et al., 2012; Wanner et al., 2016). This nomenclature distinguishes protoglomerular groups referred to as medio-dorsal (mdG; six protoglomeruli in our dataset), medio-anterior (maG; one protoglomerulus), dorsal (dG; one protoglomerulus), dorso-lateral (dlG; one protoglomerulus), lateral (lG; three protoglomeruli), ventro-posterior (vpG; two protoglomeruli), ventro-anterior (vaG; one protoglomerulus), and ventro-medial (vmG; one protoglomerulus) (Fig. 4A). Consistent with previous observations, protoglomeruli varied substantially across groups in size and shape.

**Figure 4:**
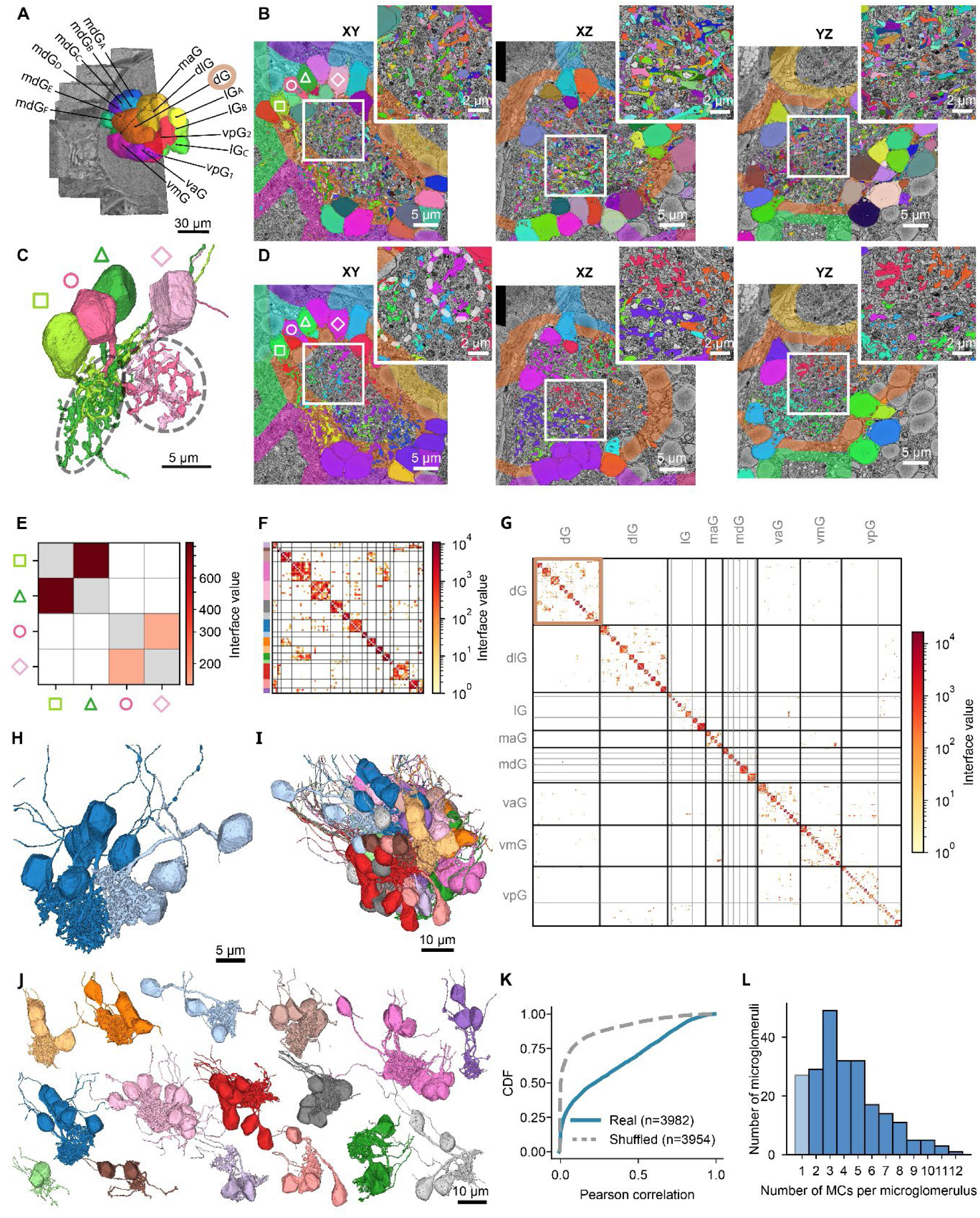
Microglomerular organization of the protoglomerular neuropil. **A.** Protoglomeruli in the left OB, delineated manually. The dG protoglomerulus is shown in more detail in following panels. Abbreviations: mdG, medio-dorsal glomerulus; maG, medio-anterior glomerulus; dlG, dorso-lateral glomerulus; lG, lateral glomerulus; vpG, ventro-posterior glomerulus; vaG, ventro-anterior glomerulus; vmG, ventro-medial glomerulus. **B.** Orthogonal views of protoglomerulus dG and adjacent protoglomeruli. MCs targeting dG are labeled in randomized colors. Four MCs within the same protoglomerulus are marked by symbols (square, circle, triangle, rhombus). A magnified view of the white-boxed region is shown in the upper-right corner of each orthogonal view. **C.** Four MCs from protoglomerulus dG highlighted in B. Neurons of different color families are part of distinct dendritic clusters (microglomeruli; dashed ellipses) within the same protoglomerulus. **D.** Same orthogonal views as in B, color-coded by dendritic cluster (microglomerulus). Dashed ellipses correspond to the clusters shown in C. **E.** Interface values of dendritic segments for the four example neurons shown in B–D. **F.** Interface values of MCs within protoglomerulus dG, sorted by cluster membership (16 clusters, manual clustering). Thin gray lines separate clusters; colored bars define the clusters shown in I. **G.** Interface values of all MCs in the left OB, sorted by protoglomerulus and cluster membership. Thick lines separate protoglomerular groups; thin lines separate individual protoglomeruli. The square marks dG (enlarged in F). Abbreviations as in A. **H.** Two adjacent microglomeruli within dG, color-coded as in F and H. Note clear separation of dendritic arbors. **I.** All microglomeruli identified in protoglomerulus dG (shown in A, B), color-coded as in F. **J.** Separated view of all microglomeruli in protoglomerulus dG (same as in I; same color code). **K.** Cumulative distribution of correlations between MC interface vectors from the real contactome and a shuffled contactome of the dG protoglomeruli (both OBs). **L.** Distribution of the number of MCs per microglomerulus, pooled across OBs. Light blue bar (1 MC) depicts solitary MCs that were not assigned to a microglomerulus and excluded from the analysis of microglomeruli.

As observed previously, most MCs had a single, compact dendritic tuft within a single protoglomerulus (Fig. 4B) (Wanner et al., 2016). We therefore assigned MCs to protoglomeruli based on the location of the centroids of their dendritic arbors (“dendritic centroids”) in each OB. Interestingly, the ventralmost mdG protoglomeruli (mdG_F_), which are closest to the midline, contained MCs with bilateral dendritic arbors innervating both mdG_F_ protoglomeruli (Supplementary Fig. 3F-H). The pair of mdG_F_ protoglomeruli was also innervated by bilaterally symmetrical dendritic arbors of four neurons with axonal projections to the telencephalon that were, however, anatomically distinct from MCs and ruffed cells (Supplementary Fig. 3E,F). As these neurons have, to our knowledge, not been described previously they were classified as “other” cells. We further observed that mdG_F_ protoglomeruli were bilaterally innervated by midline-crossing axons of OSNs (Supplementary Fig. 3G). To our knowledge, such an association between mirror-symmetrical glomeruli via midline-crossing OSN axons and MC dendrites has not been described previously.

Within the protoglomerular neuropil, neurites of different MCs were intermingled but not randomly distributed. Rather, dendritic arbors occupied distinct sub-protoglomerular volumes that overlapped extensively among small cohorts of MCs, as revealed by simultaneous visualizations of multiple MCs (Fig. 4C,D). While dendritic volumes of different MC cohorts were typically well-separated, the corresponding somata were often interspersed outside the protoglomerulus (Fig. 4D). Protoglomeruli therefore contain smaller neuropil units defined by clustered dendritic arbors of MCs that cannot be delineated by conventional histological methods or by sparse neuronal reconstructions (Fig. 4 C,D). As these structures have typical characteristics of glomeruli we refer to them as *microglomeruli* (Fig. 4 C,D,H).

To further analyze the microglomerular organization of the protoglomerular neuropil we determined the number of directly adjacent dendritic voxels (dendritic interface) between each pair of MCs (Fig. 4E), resulting in a dendritic contactome (matrix of dendritic interface values; Fig. 4G). Within this matrix, MCs were sorted first by protoglomerulus and then by clustering based on their dendritic interface. The sorted matrix showed a distinct block-diagonal structure (Fig. 4F, G), indicating strong anatomical interactions among MCs of the same cluster and weak anatomical interactions across clusters.

To explore whether the observed clusters in the interface matrix could emerge by chance, we extracted a vector for each MC from the contactome that quantified the interface area to all other MCs and compared the pairwise Pearson correlations between these interface vectors. Correlation coefficients were broadly distributed, including both low and high values. After shuffling of the contactome, high correlation coefficients were effectively abolished (Fig 4K), indicating that dendritic clustering of MCs did not emerge by chance.

To confirm that dendritic clusters of MCs correspond to microglomeruli we visualized all 16 MC clusters of protoglomerulus dG (Fig. 4F,I,J). Dendritic arbors of MCs clearly overlapped within but not between clusters, confirming that clusters in the contactome identify microglomeruli. By determining the total number of clusters in the contactome we identified 198 putative microglomeruli in total, 99 in each OB, with up to 12 MCs per microglomerulus (median: 4.0). In addition, we found 27 solitary MCs that were not associated with an identified microglomerulus but may be part of additional microglomeruli containing further, non-reconstructed MCs (Fig. 4L). The number of detecte dmicroglomeruli therefore likely underestimates the total number of microglomeruli. The number of microglomeruli per protoglomerulus was inversely related to the number of MCs per microglomerulus (Supplementary Fig. 3I).

Clusters of MCs representing microglomeruli were prominent in protoglomeruli of the dG, dlG, maG, vaG, vmG and lG groups, which segregate into multiple glomeruli at later developmental stages. Protoglomeruli of the mdG group, in contrast, were not compartmentalized into microglomeruli and do not differentiate by segregation. The two large and morphologically unique vpG glomeruli, which do also not segregate, contained multiple MC clusters that were smaller than the microglomeruli within other protoglomeruli of similar size (Braubach et al., 2013) (Supplementary Fig. 3I).

### Functional similarity of mitral cells within and across microglomeruli

In different vertebrate species, sister MCs associated with the same glomerulus show correlated spontaneous and odor-evoked activity (Chen et al., 2009; Dhawale et al., 2010). To examine whether signatures of the microglomerular organization are detectable in the activity of MCs in larval zebrafish we analyzed activity correlations between reconstructed MCs located in activity planes (n = 123) as a function of their association with micro- and protoglomeruli (Fig. 5A-D). For each pair of MCs we computed two measures representing different features of neuronal activity: (1) the Pearson correlation between odor tuning curves (signal correlation), which were constructed by averaging activity during a 2 s window covering the peak of the odor response, and (2) the temporal correlation, which was quantified by the mean Pearson correlation between unbinned spontaneous activity. Time series of spontaneous activity were constructed for each neuron by concatenating 6.7 s of activity (50 frames) prior to each odor stimulus. Similarities were then compared between pairs of MCs associated with the same or different micro- or protoglomeruli and across the two OBs.

**Figure 5:**
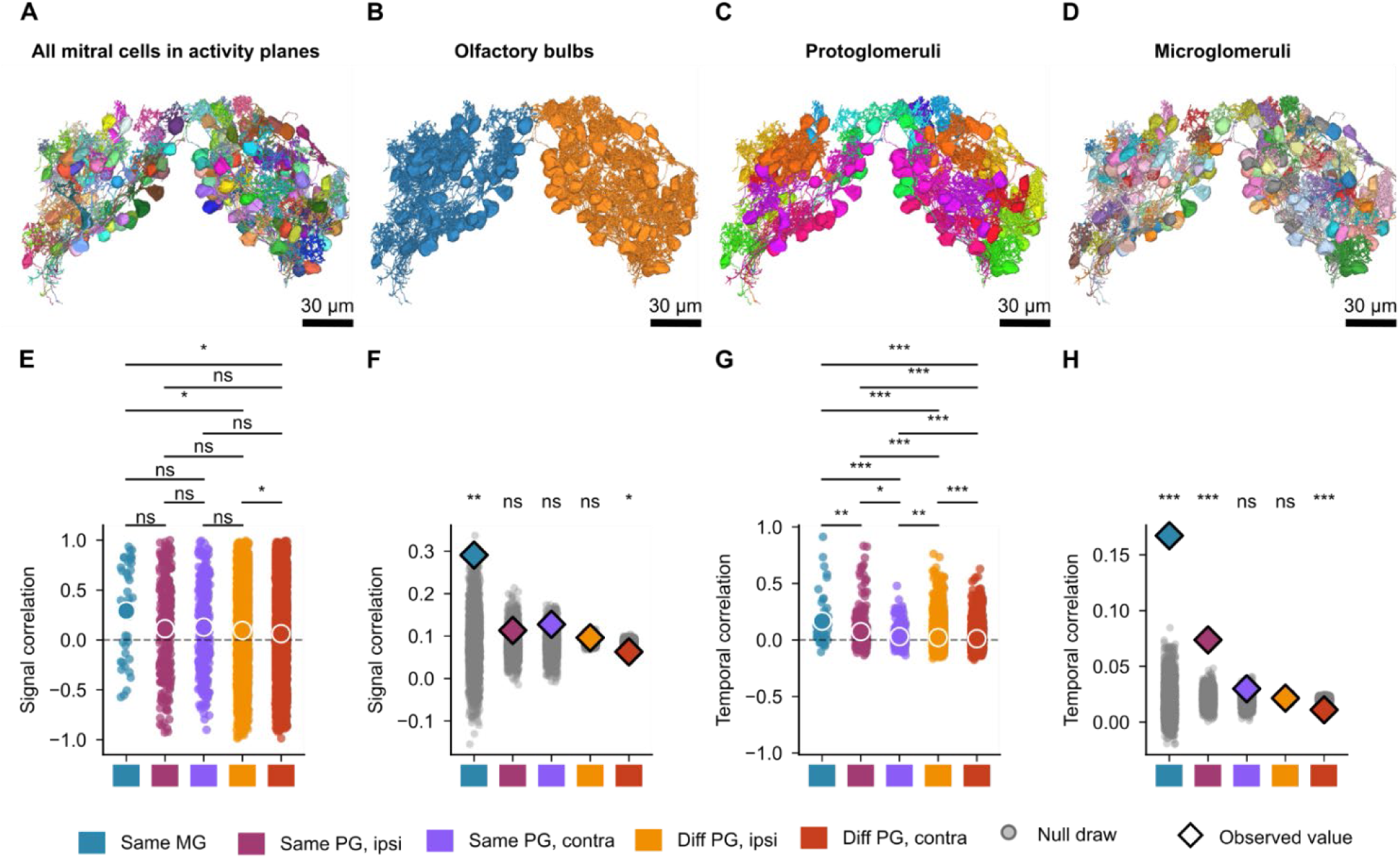
Correlated activity of MCs associated with common microglomeruli. A. MCs contained in activity planes, colored randomly. **B.** Same MCs as in A, colored by OB. **C.** Same MCs as in A, colored by protoglomerulus. **D.** Same MCs as in A, colored by microglomerulus. **E.** Tuning curve (signal) correlation for MC pairs, grouped by anatomical relationship: same microglomerulus (MG); same protoglomerulus (PG), same OB (ipsi); different OB (contra). *: P < 0.05; **: P < 0.01; ***: P < 0.001 (Mann-Whitney U test, Benjamini-Hochberg corrected). **F.** Gray: signal correlation values from E under a permutation null model, generated by jointly reshuffling MG, PG, and hemisphere labels across cells (gray points, 2,000 permutations). Colored diamonds show the observed mean correlation without shuffling for each category. **G.** Same as E, for temporal correlation. **H.** Same as F, for temporal correlation.

We found that signal correlation was higher between MCs associated with the same microglomerulus than between other MCs (Fig. 5E). However, this observation was statistically significant only across protoglomeruli, possibly because the number of MC pairs that could be sampled from both the same activity planes and the same microglomerulus was small. Moreover, a substantial fraction of MC tuning curves was not very informative about odor selectivity because MCs showed only small or no responses to any of the odors included in our panel. We therefore also used a bootstrapping approach to determine whether an observed similarity value was significantly different from an expectation based on shuffled tuning curves (Fig. 5F). This analysis showed that the observed tuning similarity was significantly higher than chance between MCs associated with the same microglomerulus, but not between MCs associated with different microglomeruli, independent of whether microglomeruli were contained within the same or different protoglomeruli (Fig. 5F). A significantly lower tuning similarity than expected by chance was found between pairs of MCs that were associated with protoglomeruli of different identity in the contralateral OB.

Highest temporal correlations between activity traces were also observed between MCs associated with the same microglomerulus (Fig. 5G). Statistically significant differences were again observed across protoglomeruli, but not within protoglomeruli, possibly because the sample size for MCs from the same microglomerulus was low. Bootstrap analysis showed that activity correlations strongly and significantly exceeded chance levels between MCs associated with the same microglomerulus (Fig. 5H). Moreover, significant correlations were found between MCs associated with different microglomeruli within the same protoglomerulus, or within the corresponding contralateral protoglomerulus. Activity correlations between MCs from protoglomeruli of different identity were indistinguishable from chance levels within the same OB and even lower than expected in the contralateral OBs. We therefore conclude that the similarity between odor tuning and temporal activity fluctuations differs between MCs depending on their association with protoglomeruli and microglomeruli, with most prominent correlations between MCs of the same microglomerulus. These results indicate that microglomeruli represent functional units within the protoglomerular neuropil.

### Microglomeruli receive distinct sensory input

We next asked whether different microglomeruli within the same protoglomerulus receive sensory input from distinct sets of OSN. To address this question we focused on protoglomerulus lG_C_ in the left OB (Fig. 6A). This protoglomerulus contained two microglomeruli, mg 146 and 193, with 9 and 11 associated MCs, respectively. We manually reconstructed 44 OSN axons within lG_C_ and identified their interface areas with reconstructed MCs. Individual OSN axons made physical contacts with 1 – 10 MCs and individual MCs were contacted by 1 – 14 OSN axons (Fig. 6B, C). While most axons terminated within lG_C_, a subset of axons passed through lG_C_ without making contacts and targeted MCs in an adjacent protoglomerulus (lG_B_, vpG_1_). All axons terminating within lG_C_ targeted MCs exclusively within lG_C_ (Fig. 6D). Hence, OSN axons innervated MCs in specific protoglomeruli.

**Figure 6:**
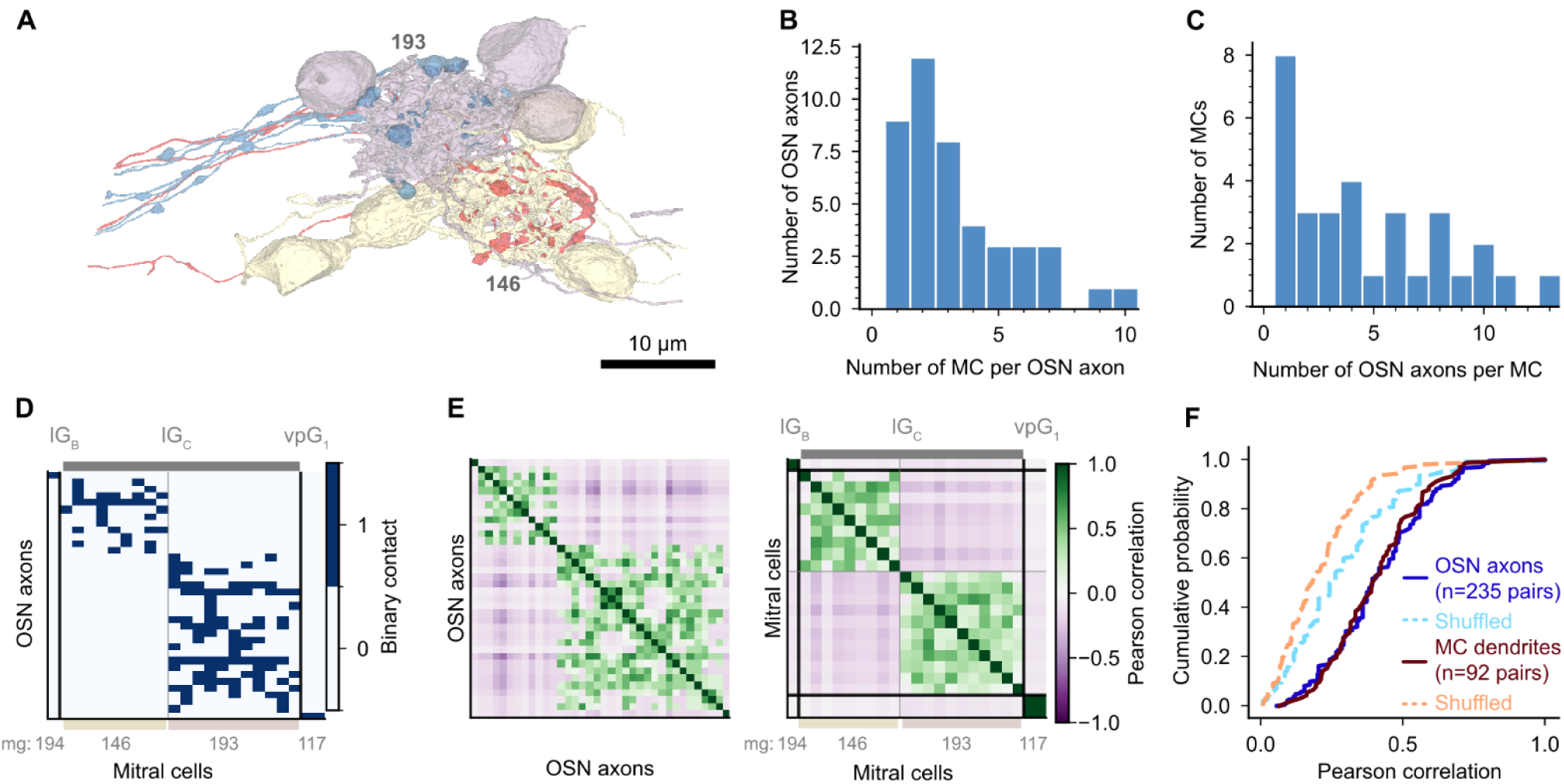
Specific targeting of microglomeruli by sensory axons. **A.** Two reconstructed microglomeruli (193, 146) and their contacting sensory axons (blue, red) in the lG_C_ protoglomerulus. **B.** Distribution of the number of sensory axons contacting each mitral cell. **C.** Distribution of the number of mitral cells contacted by each sensory axon. **D.** Binary contactome of sensory axons (rows) and MCs (columns) in protoglomerulus lG3 of the left OB. Thin gray lines separate microglomeruli. Protoglomerulus labels (top) and microglomerular assignment (mg 194, 146, 193, 117) are indicated along the x-axis. **E.** Left: Pearson correlation matrix of sensory axon interface vectors, ordered as in D. Right: Pearson correlation matrix of mitral cell interface vectors, ordered as in D. **F.** Cumulative distribution of pairwise Pearson correlation coefficients for sensory axon and mitral cell interface vectors from the real contactome and shuffled contactomes (dashed).

Within lG_C_, individual OSN axons contacted multiple MCs associated with either mg 146 or mg 193, but never MCs from both microglomeruli (Fig. 6D). This observation was confirmed by clustering OSN axons based on correlations between their interface vectors (Fig. 6E), as distinct clusters of OSNs corresponded to subsets of OSNs innervating different microglomeruli. Likewise, distinct clusters corresponding to microglomeruli were observed when the OSN-MC contactome was clustered based on correlations between MC interface vectors (Fig. 6E). Degree-preserving shuffling of the OSN-MC contactome abolished clusters, as seen by the loss of high correlation coefficients (Fig. 6F; Kolmogorov-Smirnov p = 1.7 × 10⁻¹⁴, Mann-Whitney U p = 3.7 × 10⁻¹⁷ for OSN interface vectors; Kolmogorov-Smirnov p = 1.2 × 10⁻¹⁴, Mann-Whitney U p = 9.5 × 10⁻¹⁸ for MC interface vectors). To confirm these results, we further reconstructed OSN axons (n = 9) innervating microglomerulus mg 1 within a different protoglomerulus of the contralateral OB (vaG of the right OB). As observed for microglomeruli in the left lG_C_, we found that none of the OSN axons made contacts with MCs associated with other microglomeruli in the right vaG (not shown). These results indicate that OSN axons target not only specific protoglomeruli but also specific microglomeruli within the protoglomerular neuropil. Hence, MCs associated with the same microglomerulus receive common sensory input that is distinct from sensory input to other microglomeruli.

## Discussion

Our results show that protoglomeruli of the developing zebrafish OB consist of distinct neuropil structures, referred to as microglomeruli, that were identified by reconstructing a large number of MCs within the same OB at ultrastructural resolution. This approach allowed us to comprehensively analyze the subcellular organization of the protoglomerular neuropil. Microglomeruli are defined by cohorts of MCs with overlapping dendritic arbors that are confined to distinct protoglomerular subcompartments and innervated by specific sets of sensory axons. These results indicate that a discrete array of sensory processing channels is established already at early developmental stages, around the time when the OB becomes responsive to odors and long before neuronal circuits become mature.

### Early formation of glomerular processing channels

Previous observations have not converged onto a coherent picture of glomerular development in vertebrates. In rodents and opossums, some studies indicate that glomeruli form during an early developmental time window (Malun and Brunjes, 1996; Treloar et al., 1999; Matsutani and Yamamoto, 2000; Blanchart et al., 2006; Nishizumi et al., 2019) while others reported the de novo appearance of glomeruli over multiple weeks of postnatal development (LaMantia and Purves, 1989; Pomeroy et al., 1990). The slow differentiation of some glomeruli by segregation from protoglomerular precursors in zebrafish supported the view that glomeruli emerge gradually (Braubach et al., 2013). However, we found that distinct microglomeruli are already present shortly after the OB becomes responsive to odors. We therefore conclude that discrete processing channels are established early in development even though they physically segregate into anatomically distinct neuropil units only at later stages.

Microglomeruli were defined by clustering of MC dendrites and found to be innervated by distinct subsets of OSN axons. Hence, microglomeruli exhibit classical hallmarks of glomeruli despite their small size, high packing density, and the absence of a glial boundary. Clustering delineated 99 microglomeruli in each OB, which is likely underestimating the total number of microglomeruli because our dataset included only approximately two thirds of all MCs. The expected total number of microglomeruli is therefore similar to the estimated number of glomeruli in the adult OB (∼140) (Braubach et al., 2012). Although additional glomerular units may arise later in development, these observations indicate that adult glomeruli are specified to a large extent already by microglomeruli at early larval stages.

As OSNs expressing classical odorant receptors project to single protoglomeruli in zebrafish larvae (Shao et al., 2017) it would be interesting to test the hypothesis that OSNs expressing the same classical odorant receptor target specific microglomeruli within a protoglomerulus. TAAR-expressing OSNs, in contrast, project to multiple protoglomeruli, ruling out a one-to-one mapping of receptors to glomeruli (Shao et al., 2017). V2Rs are expressed by microvillous OSNs projecting to an anatomically complex ventro-lateral region of the OB (Sato et al., 2005; Yoshihara, 2009; Braubach et al., 2012; Miyasaka et al., 2013) but their glomerular convergence has not been mapped systematically. Further studies are therefore needed to understand the mapping between chemosensory receptors, microglomeruli in the developing OB, and mature glomeruli in the adult OB.

Previous light microscopic analyses showed that ∼80% of the glomeruli in the adult OB differentiate by segregation from protoglomeruli (Braubach et al., 2013). We found that protoglomeruli undergoing segregation were compartmentalized into multiple distinct microglomeruli whereas protoglomeruli of the mdG group, which persist throughout development, were not compartmentalized. These observations further support the notion that microglomeruli are precursors of mature glomeruli in the adult OB. In addition, the OB contains two large and morphologically unique vpG protoglomeruli that do not differentiate by segregation (Braubach et al., 2013) but contain multiple small mitral cell clusters, indicating that protoglomerular subcompartments do not always give rise to distinct glomeruli. To our knowledge, no ligands or potential behavioral roles of vpG glomeruli have been identified, raising the possibility that their unique anatomical organization is associated with specific physiological functions.

The comprehensive ultrastructural analysis of MCs resulted in addition anatomical observations, including the finding that ruffed cells exist already at larval stages. Moreover, we found a specific pair of protoglomeruli (mdG_F_) that was linked across the midline through bilaterally projecting OSN axons and dendritic arbors of MCs, as well as a novel cell type. Bilaterally projecting OSN axons exist in the insect antennal lobe (Stocker, 1994) and previous studies described topographically organized axonal projections between the OBs through the anterior commissure of the telencephalon (Kermen et al., 2020). However, bilateral dendrites of MCs and other OB neurons have, to our knowledge, not been described. The pair of mdG_F_ protoglomeruli may therefore function as one unit that allows output neurons to integrate sensory inputs across a larger number of idiotypic OSNs, possibly enhancing the signal to noise ratio. The bilateral linkage of mirror-symmetrical glomeruli may occur in mdG_F_ but not other protoglomeruli because mdG_F_ protoglomeruli are closest to the midline. Alternatively, mdG_F_ protoglomeruli may subserve specific biological functions that do not require bilateral odor processing.

### Potential functions of microglomerular patterning

Individual glomeruli are functional units that integrate activity across functionally equivalent OSNs by convergence of axons onto common MCs. This integration results in fundamental computations including noise suppression and a dynamic range expansion by mechanisms involving divisive normalization (Zhu et al., 2013; Banerjee et al., 2015; Shen et al., 2025). The organization of the protoglomerular neuropil into distinct microglomeruli may therefore enhance the sensitivity, stability and concentration-invariance of odor representations in the developing OB.

Glomeruli establish an array of processing channels that are the basis for the combinatorial representation and processing of odor information. Computations in the OB include a decorrelation of activity patterns evoked by similar odorants that supports pattern classification and discrimination learning. Detailed structural and functional analyses in larval zebrafish revealed that pattern decorrelation is achieved, at least in part, by the input-specific suppression of activity among distinct cohorts of MC that respond strongly to similar odorants (Wanner and Friedrich, 2020; Friedrich and Wanner, 2021). This suppression is mediated by reciprocal connectivity between MCs of the same functional cohort and common inhibitory interneurons. Our finding that MC activity is more highly correlated within than across microglomeruli suggest that cohorts of functionally related MCs correspond to individual microglomeruli or sets of microglomeruli with similar tuning. The microglomerular organization of the developing OB may therefore provide an anatomical basis for specific higher-order connectivity underlying the decorrelation of odor-evoked activity patterns.

Generally, computations in the OB depend on specific connectivity between neurons within the same processing channel or across specific combinations of processing channels. The circuitry of the larval zebrafish OB contains already ∼50% of the MCs in the adult OB but ∼98% of interneurons, particularly granule cells, are added only at later stages (Wanner et al., 2016). Hence, the array of microglomeruli may provide a scaffold to establish a core circuitry around the time of hatching when the olfactory system becomes functional and larvae begin to express odor-dependent behaviors. Additional interneurons may then be integrated gradually into this core circuitry to refine and expand circuit function while maintaining computational core functions. It may be hypothesized that the core circuitry mediates functions of general relevance while the subsequent addition of circuit components modifies information processing based on an individual’s cumulative experience. Consistent with this hypothesis, odor exposure influenced the differentiation of glomeruli at late larval or juvenile stages but not the initial formation of protoglomeruli (Braubach et al., 2013). The microglomerular organization of the larval OB may therefore provide an anatomical basis to reconcile stability and plasticity during the lifelong development of the OB.

## Methods

### 1. In vivo functional imaging and stimulation

#### Animal husbandry and lines

Zebrafish (Danio rerio) were maintained under standard laboratory conditions on a 14:10-hour light/dark cycle at 27.5°C. Adult fish were housed in 3.5 L tanks (Tecniplast ZB30TK) equipped with continuous water circulation. Experiments were performed using Tg(α-tubulin:GCaMP6s;gad1b:DsRed) fish of both sexes in a nacre background. Fish were generated by crossing Tg(α-tubulin:GCaMP6s) (Diaz-Verdugo et al., 2019) and Tg(gad1b:DsRed) (Satou et al., 2013) lines. All experimental procedures were approved by the Veterinary Department of the Canton Basel-Stadt (Switzerland).

Embryos were raised in embryo medium (E3) containing 5 mM NaCl, 0.17 mM KCl, 0.33 mM CaCl2, and 0.33 mM MgSO4. Medium was exchanged daily. At 2 dpf, larvae were screened for expression of both fluorophores, transferred to 6-well plates, and subsequently maintained in E3 at a density of 5 larvae per well.

#### Larval preparation and mounting

Larvae were paralyzed using mivacurium chloride as described (Brustein et al., 2003; Li et al., 2005; Mack-Bucher et al., 2007) and embedded in 1.5% low-melting agarose within a custom-made silicon perfusion chamber. The body axis was inclined slightly head-up to improve optical access to the developing olfactory bulbs. Agarose covering the nose was removed to allow unobstructed odor flow through the nostrils.

#### In vivo multiplane two-photon calcium imaging

Activity measurements were then performed at 130 hpf larva measuring approximately 4 mm in body length.

We acquired GCaMP6s fluorescence images from both olfactory bulbs of a larva at 130 hpf (∼4 mm body length) using a custom-built multi-plane two-photon microscope with a remote focusing designed around a MOM body (Sutter Instruments) (Rupprecht et al., 2016). GCaMP6s was excited using a Mai Tai eHP Deep See (Spectra-physics) at 930 nm and power was modulated as a function of imaging depth by a Pockels cell (350-80LA, Conoptics). Excitation power was typically 30 mW in planes closest to the objective (20x. NA 1.0; Zeiss) and increased to 34 mW in the deepest plane (∼100 µm deeper). Remote focusing was performed using a voice-coil actuator as described (Rupprecht et al., 2016), which allowed us to acquire image volumes containing seven optical planes (256 × 512 pixels) with a field of view of approximately 205 x 205 µm^2^ at a volume rate of 7.5 Hz without physically moving the objective. As a consequence of remote focusing, one optical plane was highly distorted and excluded from further analysis. The remaining six planes were slightly tilted relative to the optical axis and their scaling varied slightly as a function of depth (Rupprecht et al., 2016). The anatomical reference stack was acquired at higher resolution (0.4 x 0.4 µm^2^ pixels, 1 µm step size in z) by moving the objective, yielding a dense set of image planes perpendicular to the optical axis. Fluorescence was detected using a GaAsP photomultiplier tube (PMT, H7422P-40MOD; Hamamatsu). Image acquisition was controlled using Scanimage software with custom modifications (Pologruto et al., 2003; Rupprecht et al., 2016).

To compensate for possible drift, a small image stack was acquired at the beginning of each experiment, spanning 9 µm around a target plane through the center of the OB with a 1.5 µm step size. During the experiment, additional stacks were acquired after each trial and possible drift was corrected by maximizing the cross-correlation to the reference stack. The anatomical reference stack for registration to the EM volume was acquired at the end of the experiment.

#### Odor stimulation

A computer-controlled system with a peristaltic pump delivered odors to the nasal epithelium via a continuous stream of E3 medium (Rupprecht and Friedrich, 2018). Flow from multiple odor channels was integrated into the main E3 perfusion line using a circularly symmetric 9-channel plastic manifold (Darwin Microfluidics Manifold 9 Ports 1/4-28 PEEK, 1/16” OD). A bubble trap (Diba Omnifit® Bubble Traps, 21940-38) was installed in the E3 delivery tube leading to the manifold to prevent air entry. The odor panel consisted of two amino acids (alanine, serine; Sigma-Aldrich), one diamine (cadaverine; Sigma-Adrich) and three bile acids (taurocholic acid [TCA], glycocholic acid [GCA], taurodeoxycholic acid [TDCA]; Sigma-Aldrich). Amino acid and bile acid stock solutions were prepared at 10 mM and 1 mM, respectively, in deionized water and stored at −20°C. Stocks were diluted 1:100 in E3 medium before each session, yielding final concentrations of 100 µM (amino acids and diamine) and 10 µM (bile acids).

Stimuli were delivered in three randomized blocks, each including one trial per odor plus two control trials (E3 exposure, spontaneous activity measurement). Stimuli were presented for 5 s, preceded by a 5 s baseline and followed by washout; inter-stimulus interval was 2–3 min. Two computer-controlled peristaltic pumps (Reglo ICC, Ismatec) ensured continuous E3 perfusion and precise odor delivery, each wheel independently driving E3 or a specific odor solution into the manifold. A custom MATLAB interface regulated pump timing and coordination.

We noticed that population activity evoked by different odors was more highly correlated in the larva selected for neuron reconstruction than in other larvae, indicating that the odor delivery system may have been contaminated by a common background odor. Analyses of odor responses were therefore limited to basic analyses of activity; quantitative analyses of pattern similarity or odor identity decoding were not performed.

### 2. Functional data preprocessing

#### Preprocessing of anatomical reference stack

To facilitate alignment of the anatomical reference stack to the EM volume the stack was first flipped horizontally along the x-axis to correct for mirroring during acquisition and then upsampled along the z-dimension by a factor of 2.5 (order-1 interpolation with reflect-mode boundary handling and anti-aliasing) to obtain isotropic voxels of size (0.4 µm)^3^. Finally, contrast was normalized using Contrast-Limited Adaptive Histogram Equalization (CLAHE).

#### LM activity planes preprocessing

Raw trial TIFFs were reshaped into 4D hyperstacks (planes × frames × y × x) and flipping horizontally as for the “anatomical reference”. Motion correction was performed using Suite2p non-rigid registration (Pachitariu et al. 2017; block size 64 × 64 pixels, maximum registration shift 0.5, SNR threshold 1.2, batch size 500) independently in each imaging plane. A stable reference image per plane was built from the final trial of the session.

#### Region of interest segmentation

Somata were segmented using Cellpose (v3.1.1.2) (Stringer et al., 2021) with a custom-trained model, applied to the plane-wise sum of registered trials. Segmentation parameters were a cell diameter of 15 µm, a cell probability threshold of 0.5, and a flow threshold of 0.0, applied to a single fluorescence channel. Detected masks were filtered by area, retaining only ROIs between 30 and 400 pixels. When segmentation was performed on individual trials the resulting masks were stitched across trials using an intersection-over-union threshold of 0.1. Segmented ROIs were manually proofread using a custom napari-based interactive tool (Sofroniew et al., 2026). Proofread masks were saved iteratively and used for all downstream trace extraction.

#### Trace extraction and normalization

Raw fluorescence traces were extracted as the mean pixel intensity within each proofread ROI mask, computed frame-by-frame for each plane and trial. Shutter-off frames (frames 0–15) were used to estimate and subtract a potential constant signal offset. Baseline fluorescence (F0) was calculated as the mean signal within a 50–100 frame window preceding stimulus onset, and dF/F was calculated as (F − F0) / F0 for each frame. Stimulus onset times were detected for each odor-delivery pump by thresholding the trial-averaged dF/F trace at the baseline mean plus 20 times the baseline standard deviation. Activity traces were then aligned to stimulus onset. dF/F traces were deconvolved into infer spiking probability by CASCADE (Rupprecht et al., 2021) using the “Global_EXC_7.5Hz_smoothing200ms_high_noise” pretrained model.

### 3. EM image acquisition and segmentation

#### Sample preparation for electron microscopy

Samples were fixed immediately after activity imaging and prepared for volume EM using the fBROPA protocol (Genoud et al., 2018). Larvae were decapitated at the head-trunk boundary at an angle such that the ventral region was slightly higher than the dorsal region when the sample was mounted vertically (anterior-side up). Samples were immersed in fixative containing 2.5% glutaraldehyde, 0.1 M cacodylate buffer, and 4% sucrose (pH 7.4). Membrane contrast was enhanced by sequential staining with formamide-reduced osmium (RedOs), osmium tetroxide, pyrogallol, and Walton’s lead aspartate at 60°C as described (Genoud et al., 2018).

After dehydration through a graded ethanol series and infiltration with an epoxy resin (Glycid ether 100, SERVA; mixed with dodecenylsuccinic anhydride and methylnadic anhydride, following (Genoud et al., 2018), the lower jaw was removed and excess resin blotted away. Effective removal of superficial resin before the subsequent embedding in conductive resin is important to minimize imaging noise at the tissue–resin–silver interface. Samples were oriented frontal-side-up and embedded in conductive silver resin (EMS EPO-TEK H20E; Electron Microscopy Science) on a metal stub to maximize conductance. Because the initial angled cut served as the mounting base, the microtome blade intersected the anterior tissue outside the OB just before cutting into the OB. This arrangement allowed us to switch from a fast approach mode to high-resolution image acquisition mode prior to cutting into the OB, thus preserving the entire OB volume.

#### Serial block-face electron microscopy and image alignment

The image stack was acquired using a Zeiss Merlin SEM equipped with an automated ultramicrotome (3View; Gatan) and a diamond knife, at [1.5 kV] and [250 pA] and a sectioning speed of 0.3 mm/s. Acquisition was managed using SBEMimage software (Titze et al., 2018). The final stack covered a volume of 231 × 245 × 115 µm^3^ at a voxel size of 10 × 10 × 25 nm^3^ (∼1 TB of data).

Raw tiles were reconstructed into a seamless volume using the Scalable Optical Flow-based Image Mesh Alignment (SOFIMA) algorithm (github.com/google-research/sofima). This procedure first stitches tiles in each horizontal section (2D) and subsequently aligns stitched cross-sections along the z-axis. Fine alignment uses flow field computation and mesh solving: the algorithm extracts image patches from a grid and compares them to reference patches to compute displacement vectors, then models the grid as a Hookean spring mesh, numerically minimizing energy to balance elastic forces preserving image structure against flow-driven forces pulling images into alignment. The aligned volume is rendered from this mesh optimization.

#### Neuronal reconstruction and segmentation

Base (fragment-level) segmentation was performed using a flood-filling network (FFN; Januszewski et al. 2018) trained on an EM dataset from the adult zebrafish brain. Prior to segmentation, raw EM images from the larval brain were domain-adapted to the training data’s staining/imaging statistics using a Segmentation-Enhanced CycleGAN (SECGAN), a CycleGAN variant that adds a segmentation-consistency constraint to prevent structural hallucination during translation (Januszewski and Jain, 2019). Fragments were agglomerated into complete neuronal reconstructions via supervised edge classification on a candidate adjacency graph, following the standard Google connectomics pipeline (Januszewski et al., 2018; Lu et al., 2021). Cellular compartment identity (soma, nucleus, dendrite, axon) was assigned using custom tools fine-tuned on manually annotated voxels from the present dataset spanning the four compartment classes. This compartment classifier’s output was used both to suppress biologically implausible agglomeration edges (e.g., axon–dendrite or cross-soma merges) and to restrict downstream connectivity analyses to relevant compartments. The agglomeration graph was maintained via a mutable change-stack and equivalence-mapping service, allowing reconstruction identities to be updated non-destructively during proofreading. Manual proofreading was performed through an interactive graph-editing interface (Mönig, 2020a)

#### Voxel contact interface quantification

Voxel-level interface values between adjacent base segments were computed as the number of shared boundary voxels between every pair of adjacent base segments across the EM volume. Base-segment interface counts were aggregated to the agglomeration level by mapping each base segment to its parent agglomeration and summing shared-voxel counts across all bridging base-segment pairs, producing the agglomeration-level interface tables (contactome).

### 4. Agglomerate classification

#### Neuronal cell-type classification

Neurons were manually classified with a classification tool into mitral cells (MC) and interneurons (IN) using a custom interactive classification viewer (Mönig, 2024). As an independent validation and extension of this classification, gad1b:DsRed expression was quantified directly from EM-registered nuclei by sampling red-channel fluorescence intensity within each nucleus mask from the anatomical light-microscopy stack (Gaussian-denoised, per-slice percentile-normalized). Because signal intensity decayed with imaging depth, a per-sample depth-correction curve (exponential decay model) was fit using manually classified mitral cells and interneurons as ground truth, and correction factors were applied to normalize the signal along the z axis. A classification threshold and a logistic (sigmoid) probability model were then fit to the depth-corrected signal by maximizing classification accuracy against ground-truth labels, with model performance quantified via ROC-AUC. This procedure was used to assign each nucleus to one of three classes with respect gad1b:DsRed expression (positive/negative/ambiguous).

#### Manually drawn region masks

OB boundaries and protoglomerular territories were delineated manually in the EM volume. Each segmented cell was assigned to a region by testing whether its representative coordinate (nucleus centroid for cell-body-based assignment, or dendrite median for protoglomerulus assignment) fell within the corresponding mask, evaluated at a voxel scale of 160 × 160 × 320 nm. Brain area assignments followed the same approach, using each agglomeration’s nucleus centroid as the query point.

Protoglomerular groups were labeled according to the nomenclature of Braubach et al. (Braubach et al., 2012). In some groups (lG, mdG) individual protoglomeruli showed a consistent, mirror-symmetrical arrangement in the two OBs of the larva analyzed here but could not be matched unequivocally to individual protoglomeruli described previously. We therefore distinguished protoglomeruli within these groups by letters (A, B,…) rather than by numbers (Braubach et al., 2012).

#### Semi-automated microglomerulus assignment

Mitral cells were assigned to microglomeruli using a nearest-prototype cosine similarity classifier built on dendrite-to-dendrite contact profiles: for each mitral cell, contact strength with every other mitral cell was quantified by the contact area taken from the symmetric agglomeration-level interface matrix (restricted to dendritic segments). Each cell’s contact vector was then L2-normalized. For every microglomerulus with existing confirmed membership, a prototype vector was computed as the mean of its member cells’ normalized contact vectors (itself renormalized), and each unassigned cell was assigned a similarity score to every prototype. The three highest-scoring candidate microglomeruli, together with the similarity margin between the top two candidates, were reported for manual visual confirmation before acceptance.

### 5. Multi-modal image registration

#### Nuclei segmentation

Nuclei in the EM anatomical stack were segmented in 3D using Cellpose (v3.1.1.2) (Stringer et al., 2021), human-in-the-loop, with filtering and removal of non-neuronal objects. Nuclei of the alpha-tubulin calcium marker of LM activity planes were segmented using the same human-in-the-loop Cellpose approach with a custom model as described in “Region of interest segmentation”.

#### LM anatomical stack to LM activity plane alignment

Registered activity planes were additionally aligned to the preprocessed LM anatomical stack per plane using tiled template matching (2 × 3 tile grid, correlation threshold 0.4, nearest-neighbor interpolation), with a similarity transform computed from a chosen reference trial and applied to all remaining trials of that plane. A custom script aligned LM planes by tiling them in feature-important areas and finding the highest image correlation in the LM stack independently for each tile. The LM stack was then rotated parallel to the plane where the best tile-centroid position was found, iterating until image correlation for each tile exceeded at least 0.6.

#### EM to LM anatomical stack alignment

Rough alignment was performed by rotating the EM stack axes and reslicing to match the dorsal view of the LM stack. Using BigWarp (Bogovic et al., 2016), coarse landmarks (blood vessels, identifiable neuropil areas) were used to align the stacks via an affine transform. To minimize manual landmark-placement error, an unmatched landmark list based on extracted centroids of previously segmented EM nuclei was used as exact landmarks. These landmarks were drawn randomly and matched iteratively.

#### EM to LM activity plane alignment

Functional ROIs were additionally mapped directly onto EM segmentation by projecting each ROI’s centroid onto an EM mask warped into the light-microscopy reference stack, using vectorized nearest-label lookup.

#### Quantitative analysis and validation of registration

To quantify the accuracy of LM–EM centroid alignment we evaluated three metrics. (1) Mean alignment error: the Euclidean distance between each transformed LM centroid and its nearest-neighbor match in the EM point cloud. (2) Neighbor distance ratio (NDR): the nearest-neighbor distance normalized by the mean distance to the second- and third-nearest neighbors in the EM point cloud (k = 2 additional neighbors), yielding a point-density-corrected confidence score. Low NDR values indicate a match that stands out clearly from local point density, whereas values approaching one indicate an ambiguous match with a competing candidate of similar distance. NDR itself was not thresholded. Points with a raw alignment error above 5 µm were excluded from downstream summary statistics. (3) A separation score, quantifying the difference between the residual distance to the assigned match and the distance to the nearest competing candidate (adapted from Bae et al. 2025).

#### Sensitivity to transformation perturbation

Registration robustness to small transformation errors was assessed by perturbing the moving point cloud with independent grid searches over rotation, translation, and anisotropic scaling, applied about the point cloud’s own centroid along each spatial axis. For each perturbation magnitude and axis, mean NDR and error score (points below the error threshold) were recomputed, yielding axis-specific sensitivity profiles.

#### Coordinate mapping and data integration

EM segment identities were assigned to functional ROI and nuclear centroids through a multi-step coordinate-to-segment pipeline. EM masks were downsampled, coarse-LM-aligned, and fine-LM-aligned into LM space. Centroids of functional ROIs and LM nuclei were then assigned to EM masks converted into EM space. Each centroid was then queried against the segmentation volume using an iterative expanding-box search (initial size 1 voxel, expanding in increments of 5 up to 50 voxels, minimum 1 valid voxel), the majority label within each search box was assigned as the matching base segment, with a reported confidence equal to the fraction of box voxels sharing that label. Stable base segments were subsequently mapped to their current agglomeration IDs via the Google BrainMaps API functions (Mönig, 2020b), and agglomerations sharing more than one base segment were flagged for manual quality-control review. Manually reviewed merge/duplicate decisions were synced back into a persistent exclusion/flag list used to filter erroneous mappings in all downstream analyses.

### 6. Functional data analysis

#### Response similarity metrics

Signal correlation was computed as the Pearson correlation between odor responses, defined as the mean activity over a 2 s response window covering the peak (frames 145–160) within the 5 s odor stimulus presentation (frames 130–168), averaged across trial repetitions for each odor. This measure quantifies pairwise similarity between the odor tuning curves of MCs. Temporal correlation was computed as the Pearson correlation between mean, trial-averaged raw activity traces during a 6.7 s pre-stimulus baseline period (frames 50–100), concatenated across trials. This measure quantifies the similarity between spontaneous temporal fluctuations in activity in the absence of odor stimulation.

#### Statistical comparison of anatomical groups

Response similarity distributions were compared using pairwise two-sided Mann-Whitney U tests between categories, with p-values corrected for multiple comparisons using the Benjamini-Hochberg procedure. To test whether each relationship category showed similarity beyond what anatomy-independent chance would predict, a joint label-permutation null was constructed by shuffling microglomerulus, protoglomerulus, and hemisphere identity together across cells (2,000 permutations) and recomputing the five-way classification under each shuffle. The observed group mean was compared against its own empirical null distribution using a two-sided permutation p-value.

### 7. Structural data analysis

#### Cell-cell connectivity matrix construction

Sensory axon–to–MC and MC–to–MC contact matrices were derived from the agglomeration-level interface tables, symmetrized by summing interface size over both id-ordering directions and restricting axon–MC pairs to strictly validated sensory-axon/mitral-cell type combinations. Weighted matrices retained summed dendrite–dendrite interface size as the edge weight, and binary matrices were derived by thresholding weighted matrices at zero contact. Sensory axon rows were ordered by their column of peak contact and descending peak strength, while MC columns were hierarchically sorted by OB, protoglomerular and microglomerular identity to reveal block-structured connectivity.

Statistical significance of observed connectivity structure was assessed against a degree-preserving null model generated with the Curveball algorithm, which iteratively swaps non-shared partners between randomly selected row pairs while preserving each sensory axon’s and mitral cell’s total number of binary partners. Because Curveball preserves only binary topology, corresponding weighted null matrices were generated by randomly redistributing the empirical pool of nonzero interface sizes onto the null matrix’s contact positions, thereby preserving the overall weight distribution while randomizing which specific MC–MC pairs carried which weight. Mixing was run for 5,000 row-pair swaps per null matrix.

#### Structural comparisons against the null model

Pearson correlation and cosine similarity were computed between all pairs of sensory axons (based on shared mitral cell targets) and between all pairs of mitral cells (based on shared sensory axon inputs). The resulting real and shuffled similarity-value distributions were compared using two-sample Kolmogorov-Smirnov and Mann-Whitney U tests.

#### Software and visualization tools

All quantitative analyses were performed in Python using NumPy and pandas for array and tabular data handling, SciPy for statistical testing (Mann-Whitney U, two-sample Kolmogorov-Smirnov, Kruskal-Wallis, Pearson correlation), and scikit-learn for cosine similarity, pairwise distance computation. Matplotlib and seaborn were used for all static data visualization, including heatmaps, CDF/histogram overlays, violin and swarm plots. Neuroglancer was used for reconstruction and visualization of EM datasets.

## Acknowledgements

We thank Nesibe Temiz, Bo Hu, Estelle Arn, Tim-Oliver Buchholz, Alessandro Motta, Oded Mayseless, and Benjamin Titze for important input, the Friedrich group for fruitful discussions, and Miguel Gomes and team for fish care. This work was supported the by Novartis Research Foundation, by the European Research Council (ERC) under the European Union’s Horizon 2020 research and innovation program (grant agreement nos. 742576, 101167289), by an EMBO postdoctocal fellowship to JK (ALTF 818-2023), and by the Swiss National Science Foundation (grants no. 31003A_172925/1, 310030_219500, 310030_212236).

---

**Supplementary Figure 1:**
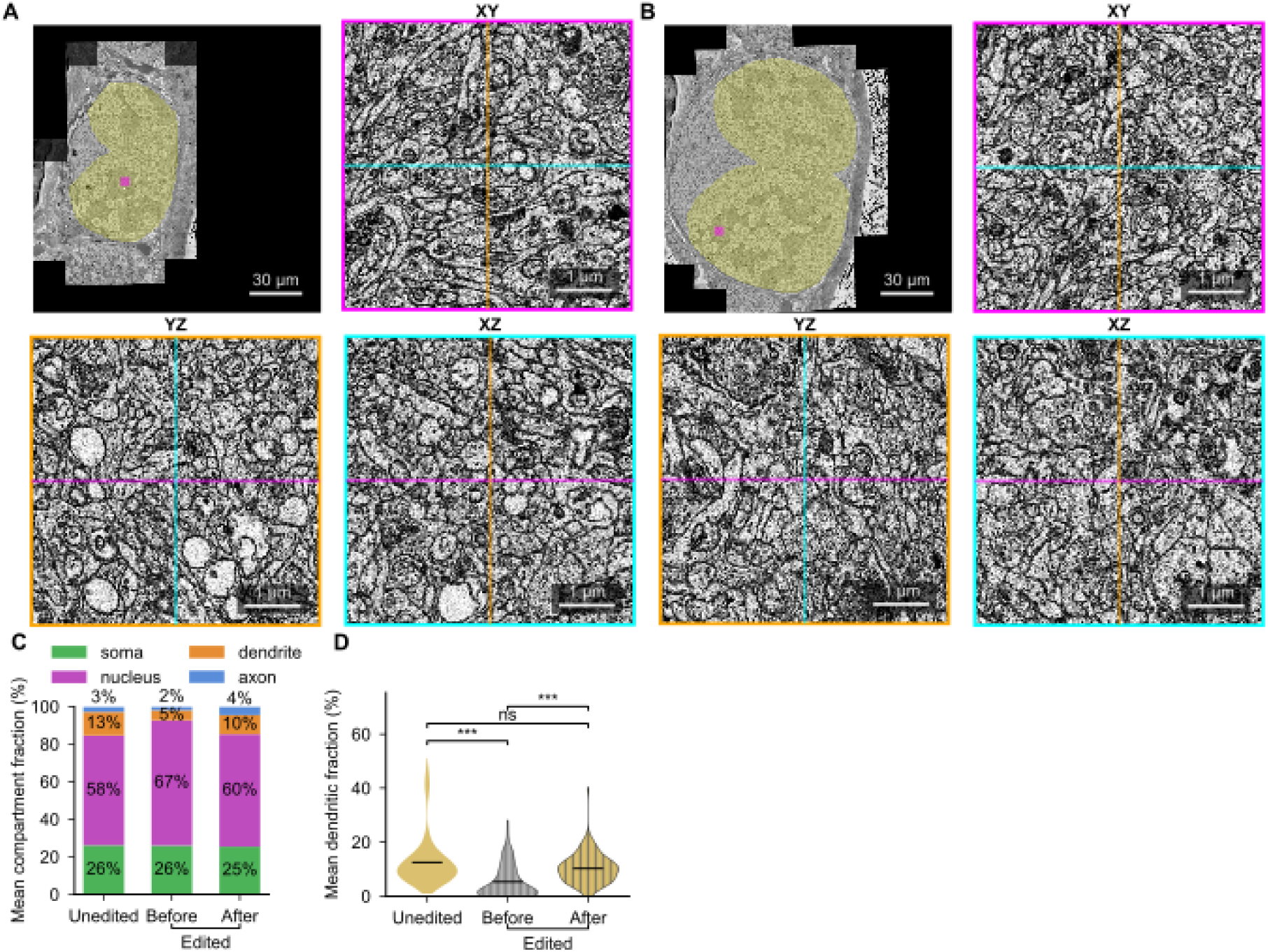
EM volume and neuron reconstruction: additional observations. **A,B.** Overview and high-resolution orthogonal cross-sections from two neuropil-rich subregions distant from the subregion shown in Fig. 2C. Note high resolution and contrast throughout the OB. **C.** Fraction of voxels assigned to different semantic categories in MCs that were not selected for proofreading (Unedited), and in MCs selected for proofreading before and after editing. Note that the fraction of dendritic voxels increased after proofreading. **D.** Fraction of voxels classified as dendritic in MCs that were not selected for proofreading (Unedited), and in MCs selected for proofreading before and after editing. Editing significantly significantly increased the fraction of dendritic voxels in MCs selected for proofreading. After editing, the fraction of dendritic voxels was comparable to that of MCs that that were not selected for proofreading, indicating that targeted proofreading resulted in more complete dendritic arbors. Statistical comparisons: Kruskal-Wallis test, ***: p < 0.001.

**Supplementary Figure 2:**
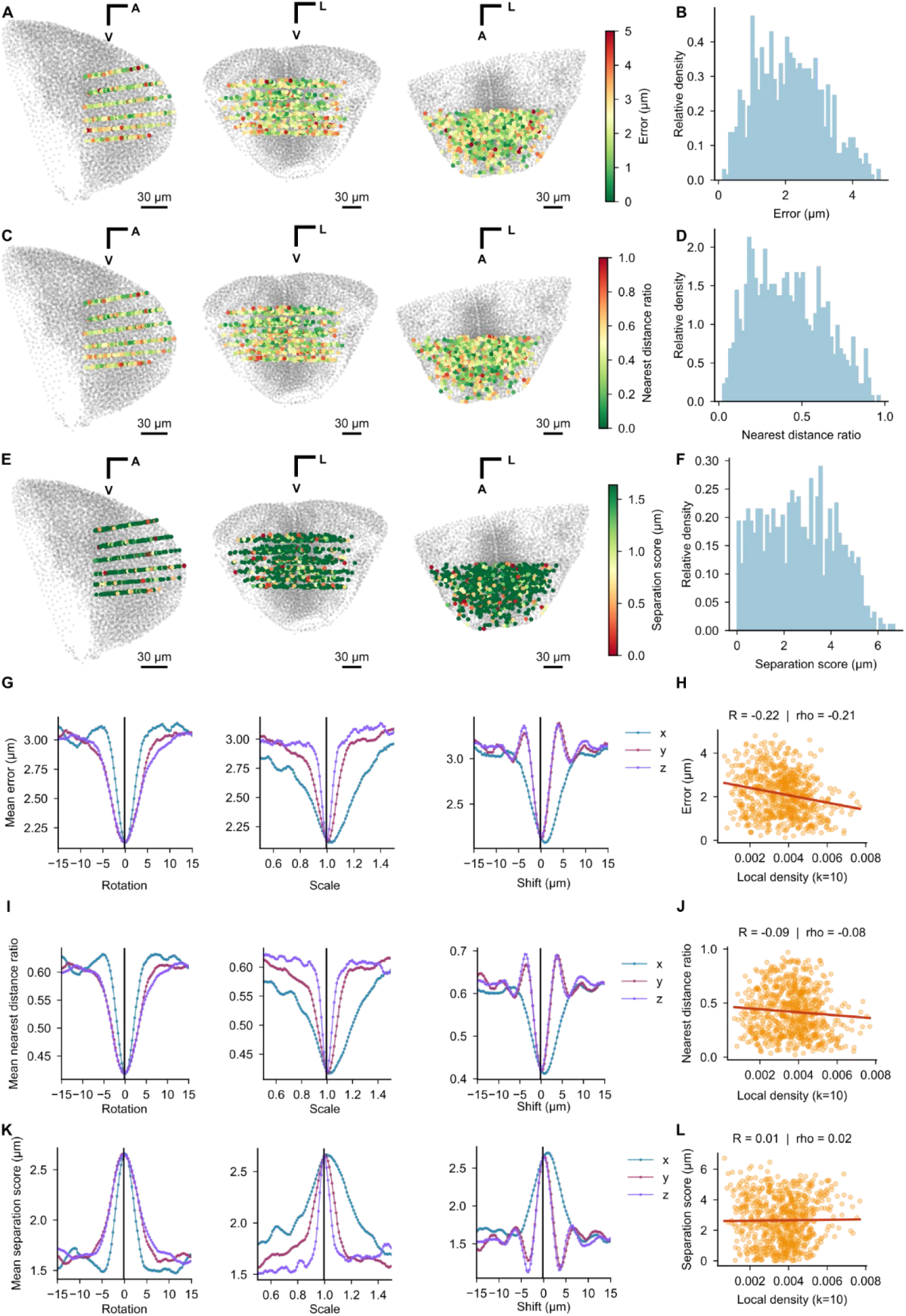
LM-EM registration: quantitative analysis. **A.** Point cloud of LM centroids in activity planes color coded by alignment error. Panels show projections along different directions as in Fig. 3D. **B.** Histogram of alignment errors across all centroids. **C.** Same as (A) for NDR. **D.** Same as (B) for NDR. **E.** Same as (A) for separation score. **F.** Same as (B) for separation score. **G.** Mean alignment error as a function of rotation, scale or shift applied to LM centroids. **H.** Alignment error as a function of and local point cloud density. **I.** Same as (G) for NDR. **J.** Same as (H) for NDR. **K.** Same as (G) for separation score. **L.** Same as (H) for separation score

**Supplementary Figure 3:**
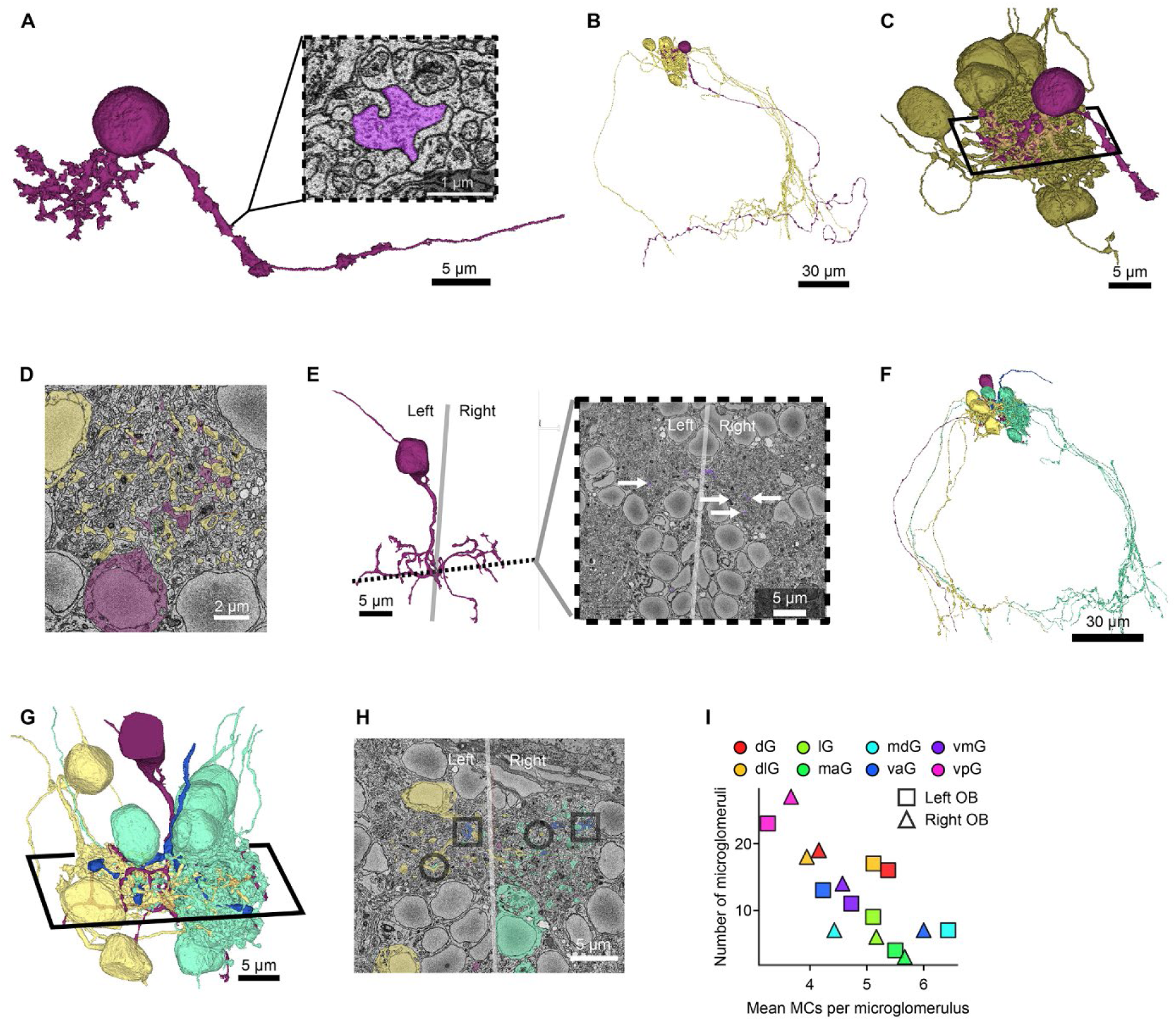
Ruffed cells and midline-crossing cells. **A.** Ruffed cell. EM image shows cross-section through the ruff with an input synapse. **B.** Same ruffed cell as in A and MCs of the same microglomerulus (yellow). Note that the dendritic arbor but not the ruff are embedded in the microglomerular neuropil. **C.** Close-up of B. Parallelogram indicates the area magnified in D. **D.** Ruffed cell (purple) and MC (yellow) dendrites in an EM cross-section through the microglomerulus shown in B and C. **E.** Neuron contacting the pair of mdGF protoglomeruli across the midline. Arrows indicate dendrites in the left and right OB. Gray line indicates midline. **F.** Same neuron as in E (magenta) shown together with other projection neurons associated with mdG_F_ protoglomeruli (yellow and green) and an OSN axon contacting both protoglomeruli (blue). Additional OSN axons with bilateral projections to both mdGF protoglomeruli were found but are not shown for simplicity. **G.** Close-up of F. Parallelogram indicates the slice shown in H. **H.** Slice through the EM volume as marked in G. Circles mark inter-bulbar connections of MCs. Squares mark OSN contacts in both OBs. **I.** Number of microglomeruli per protoglomerulus (y-axis) as a function of the mean number of mitral cells per microglomerulus within the same protoglomerulus (x-axis). Squares and triangles represent the left and right OB, respectively; symbols are color-coded as in A for both OBs. Abbreviations as in A.

## Notes

### Competing Interest Statement

The authors have declared no competing interest.

